# Direct estimation of HDL-mediated cholesterol efflux capacity from serum

**DOI:** 10.1101/396929

**Authors:** Sanna Kuusisto, Michael V. Holmes, Pauli Ohukainen, Antti J. Kangas, Mari Karsikas, Mika Tiainen, Markus Perola, Veikko Salomaa, Johannes Kettunen, Mika Ala-Korpela

## Abstract

High-density lipoprotein mediated cholesterol efflux capacity (HDL-CEC) is a functional attribute that may have a protective role in atherogenesis. However, the estimation of HDL-CEC is based on *in vitro* cell assays that are laborious and hamper large-scale phenotyping. Here, we present a cost-effective high-throughput nuclear magnetic resonance (NMR) spectroscopy method to estimate HDL-CEC directly from serum. We applied the new method in a population-based study of 7,603 individuals including 574 who developed incident coronary heart disease (CHD) during 15 years of follow-up, making this the largest quantitative study for HDL-CEC. As estimated by NMR-spectroscopy, a 1-SD higher HDL-CEC was associated with a lower risk of incident CHD (hazards ratio 0.86; 95%CI 0.79-0.93, adjusted for traditional risk factors and HDL-C). These findings are consistent with published associations based on *in vitro* cell assays. These corroborative large-scale findings provide further support for a potential protective role of HDL-CEC in CHD, and substantiate this new method and its future applications.

## INTRODUCTION

Circulating high-density lipoprotein (HDL) particles mediate reverse cholesterol transport by carrying excess cholesterol from the periphery, such as the arterial wall, to the liver for excretion into the bile^1^. HDL cholesterol (HDL-C) is an established epidemiological risk factor for cardiometabolic conditions^2^. However, the role of HDL-C remains unclear since most HDL-C increasing therapies have, on the whole, failed to prevent cardiovascular events^3^, and Mendelian randomization studies have given consistent evidence that HDL-C is not causal in the development of cardiovascular disease^4^. Even though the recent REVEAL trial^5^, of the cholesteryl ester transfer protein inhibitor anacetrapib, resulted in a lower risk of major coronary events, rather than providing evidence for a causal role of HDL-C, these findings were entirely proportional to the reduction in apolipoprotein B-containing lipoproteins^6,7^.

Therefore, the totality of evidence does not support a causal role for HDL-C in CHD. This has shifted the focus of HDL research from circulating HDL-C concentrations to other aspects of HDL, such as the functional attributes of HDL particles^1,8^. Cholesterol efflux capacity of HDL (HDL-CEC), which quantifies the ability of HDL particles to extract cholesterol from lipid-laden cells, has emerged as the most widely used metric for HDL function. HDL-CEC reflects the combined action of various HDL particles via multiple cellular pathways^9^: intracellular cholesterol is extracted by HDL via adenosine triphosphate (ATP)-binding cassette (ABC) transporters (ABCA1 and ABCG1), scavenger receptor B1 (SR-B1) and simply by passive diffusion^9^. There are multiple cellular efflux assays available either to target a specific pathway or their combination^8,10,11^. The most common assay to analyse HDL-CEC uses cyclic adenosine monophosphate (cAMP)-treated J774 murine macrophages with radiolabelled cholesterol^10,12–14^. HDL-CEC measured in cAMP-treated J774 cells incorporates all the abovementioned pathways^12^. A fluorescence-labelled cholesterol method has also been used^11,14^. Despite variation in the relative contributions from the different efflux pathways, the correlation between HDL-CEC quantified by these two assays is quite high ^11,15^.

In recent years, several studies have investigated the association of HDL-CEC with cardiovascular risk in individuals, with quantification of HDL-CEC mainly by cAMP-treated J774 cells^10,11,14,16^. These studies were recently summarised in a meta-analysis that strengthened the evidence that HDL-CEC is inversely associated with cardiovascular risk, with the association being independent of HDL-C concentrations^14^. However, results from individual studies were inconsistent^14^ and large-scale evidence on HDL-CEC and cardiovascular outcomes is currently limited to two studies, one by Rohatgi et al^11^ and the other by Saleheen et al ^10^. Both of these studies identified inverse associations between HDL-CEC and cardiovascular events independent of established cardiovascular risk factors, including HDL-C and/or apolipoprotein A1 (apoA1). Interestingly, a recent study suggested HDL-CEC to be heritable, independently of HDL-C^17^. In epidemiological studies, HDL-CEC is associated moderately with HDL-related parameters, such as HDL-C^10,11,17^, HDL size^11,17^ and total HDL particle concentrations (HDL-P)^11,17^, but weakly with clinical variables (such as BMI and blood pressure)^10,11^. Large-scale characterisation of the associations of HDL-CEC with multiple cardiometabolic risk factors, as well as HDL subclasses, is currently lacking. The relative paucity of large-scale epidemiology is most likely due to the complexity and cost of cellular HDL-CEC assays. Novel approaches are needed to facilitate such measurements and enable large-scale investigations of the epidemiological role, genetic architecture, and potential causality of HDL-CEC.

To this end, we have developed a high-throughput cost-effective alternative approach to the estimation of HDL-CEC through serum nuclear magnetic resonance (NMR) spectroscopy. Recent advancements in experimentation and automated molecular quantifications have made applications of quantitative NMR in epidemiology and genetics increasingly common in recent years^18,19^. These advances have taken NMR-based approaches into large-scale research beyond their well-known role in detailed quantification of lipoprotein subclasses, particles and lipids^18–20^. We show here that it is possible to estimate HDL-CEC from serum NMR spectra and that quantification recapitulates the characteristics of *in vitro* HDL-CEC in cAMP-treated J774 cells. This report presents the new high-throughput methodology and confirmatory results regarding the associations of HDL-CEC and incident CHD in a large-scale prospective epidemiological study.

## RESULTS

### HDL-CEC estimation via serum NMR spectroscopy

An overview of the study is described in **Supplementary Fig. 1**. Using a Bayesian linear regression model, we established a quantitative relationship between the NMR spectral regions of lipoprotein resonances and *in vitro* measured HDL-CEC. We found a good correspondence (R^2^ of 0.83) between NMR-based HDL-CEC and *in vitro* HDL-CEC (online Methods and **Supplementary Fig. 2**).

**Figure 1:**
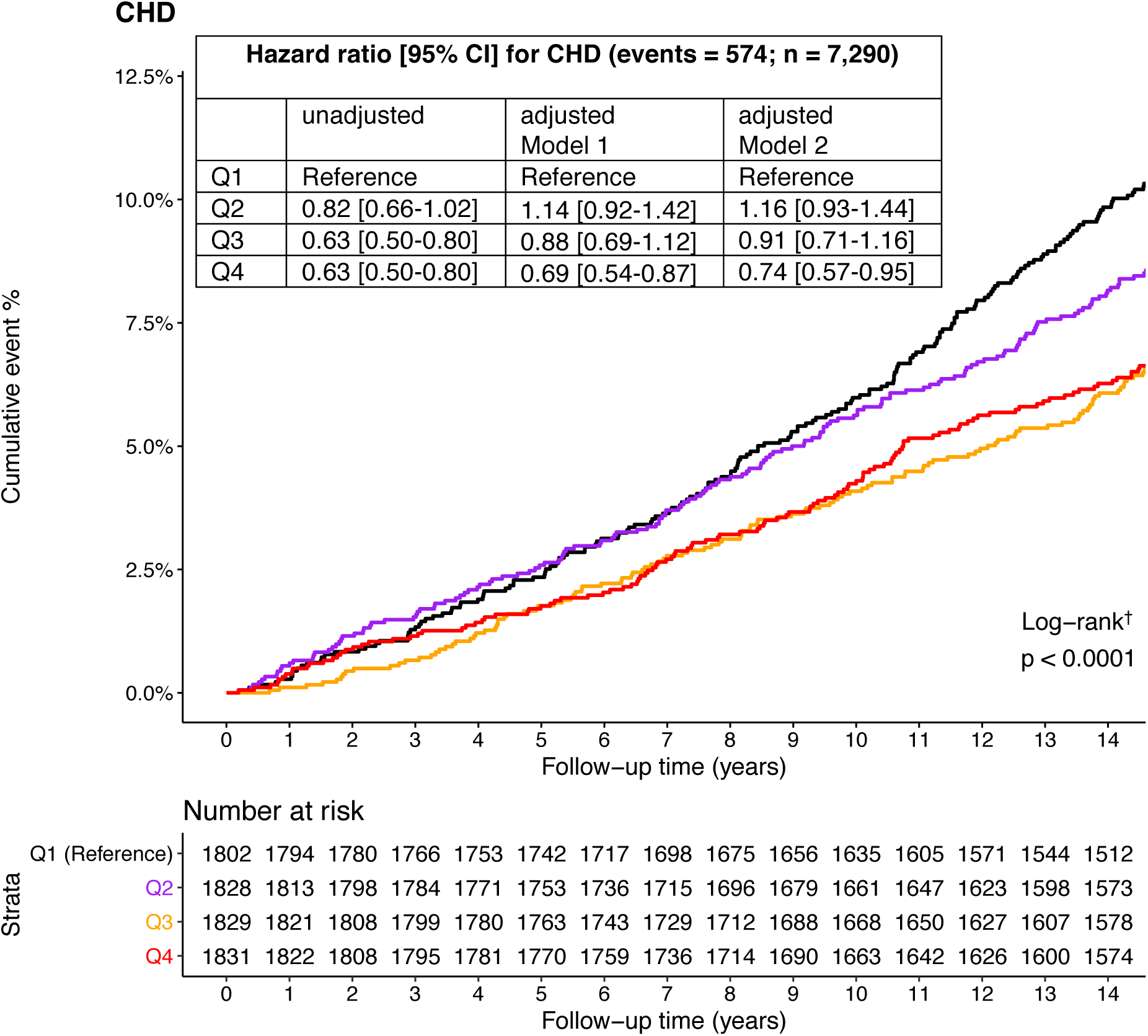
Kaplan-Meier cumulative incidence and hazard ratios of CHD by quartiles of NMR-based HDL-CEC. Hazard ratios were calculated by Cox proportional hazard models with the lowest quartile as the reference group. HDL-C; high-density lipoprotein cholesterol concentration, apoAl; apolipoprotein A1 concentration, HDL-P; total high-density lipoprotein particle concentration (a sum of the individual HDL subclass particle concentrations), TG; triglycerides. Model 1: Traditional risk factors (age, sex, geographical region, diabetes, mean arterial blood pressure, blood pressure treatment, smoking, log BMI, total cholesterol, log TG, lipid lowering treatment), HDL-C Model 2: Model 1, apoAl and HDL-P † Additional statistics for Log-rank test (chi-squared = 22.2, df = 3,two-sided)

**Figure 2:**
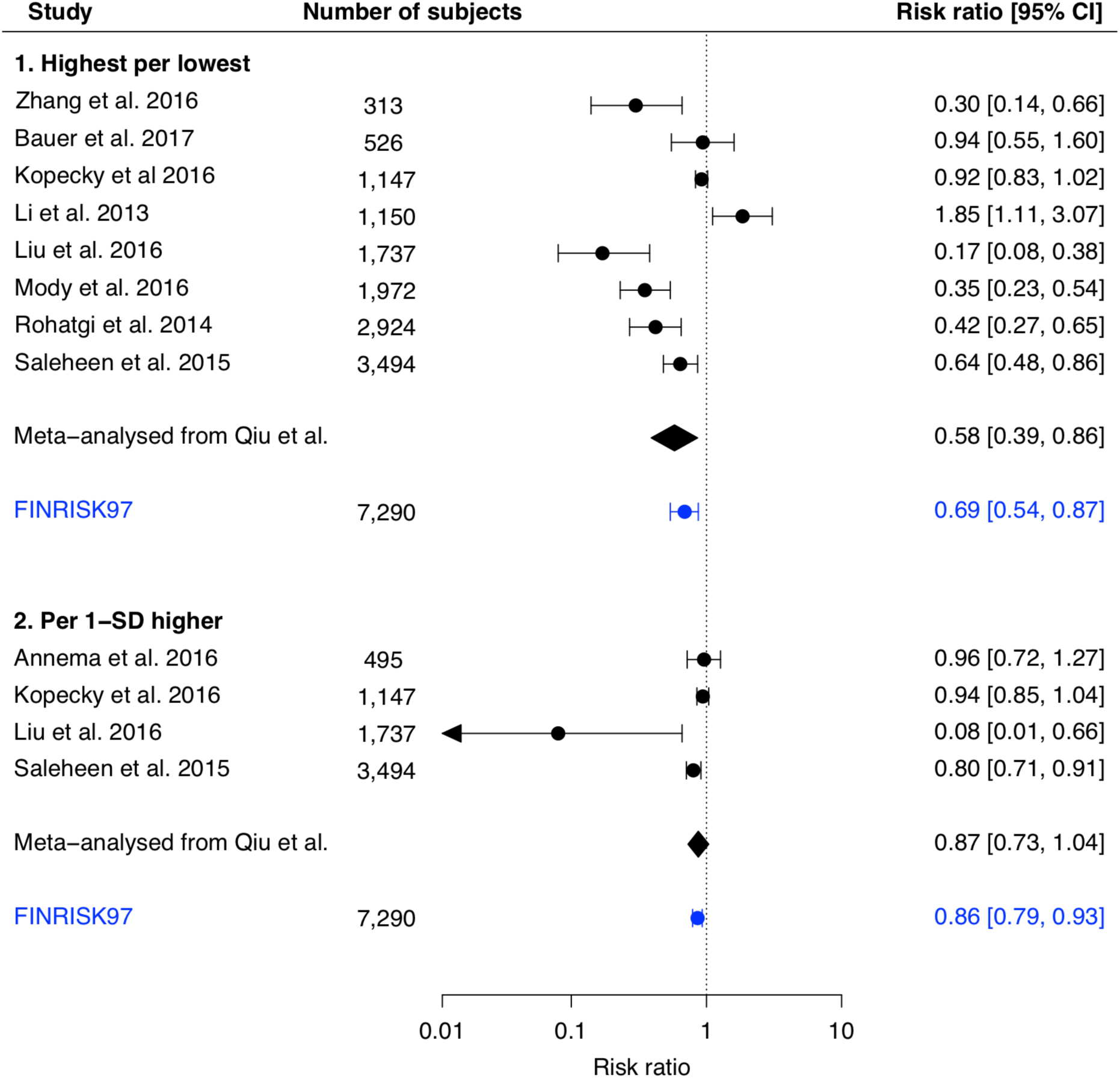
Associations of HDL-CEC with cardiovascular risk in the FINRISK97 (NMR-based) compared to previous studies (*in vitro* cellular assays). Previous studies (black colour); risk ratios (RR) and their meta-analysed results were taken from a recent meta-analysis^14^ investigating the association of HDL-CEC with cardiovascular outcomes. Highest vs lowest denote the RR for CVD comparing the highest to the lowest HDL-CEC quantiles defined in each study. The results in the current FINRISK97 cohort study (in blue); RR was adjusted for age, sex, geographical region, diabetes, mean arterial blood pressure, blood pressure treatment, smoking, log BMI, total cholesterol, log TG, lipid lowering treatment and HDL-C. Highest vs lowest refers to top vs bottom quartile. There was no evidence of heterogeneity between the current study estimate and the pooled estimate from the meta-analysis by Cochran’s Q test, two-sided (P value = 0.46, Q = 0.54, df = 1) and P value = 0.91, Q = 0.01, df = 1) for highest vs lowest and per-l-SD higher, respectively.

### Association of HDL-CEC with incident CHD and CVD events

Several studies have found inverse associations between HDL-CEC and cardiovascular outcomes, although results are heterogeneous. We were interested to see whether the analysis using NMR-quantified HDL-CEC would replicate findings by prior studies based on *in vitro* HDL-CEC assays. We analysed the association of NMR-based HDL-CEC with incident cardiovascular events in a large-scale population-based FINRISK97 cohort (7,290 individuals with 574 incident CHD events and 7,231 individuals with 789 incident CVD events during 15 years of follow-up, online Methods). The Kaplan-Meier curves in **Fig. 1** illustrate the association of NMR-based HDL-CEC quartiles with risk of incident CHD events during follow-up for the individuals in the FINRISK97 cohort. There was a dose-response inverse association between increasing quartiles of HDL-CEC and risk of CHD; for the top vs. bottom quartile of NMR-based HDL-CEC, the HR was 0.63 (95% CI 0.50-0.80) (**Fig. 1**). Detailed analyses with multiple adjustments are presented in **Supplementary Table 1**. A similar association was obtained for the association of HDL-CEC with CVD (**Supplementary Fig. 3**). HDL-CEC remained associated with CHD and CVD events even after adjustment for traditional risk factors and all other HDL-related measures, including HDL-C, apoA1 and total HDL-P; top vs. bottom quartile HR for CHD 0.74 (95% CI 0.57-0.95) (**Fig. 1**) and HR for CVD 0.79 (95% CI 0.64-0.98) (**Supplementary Fig. 3**).

**Figure 3:**
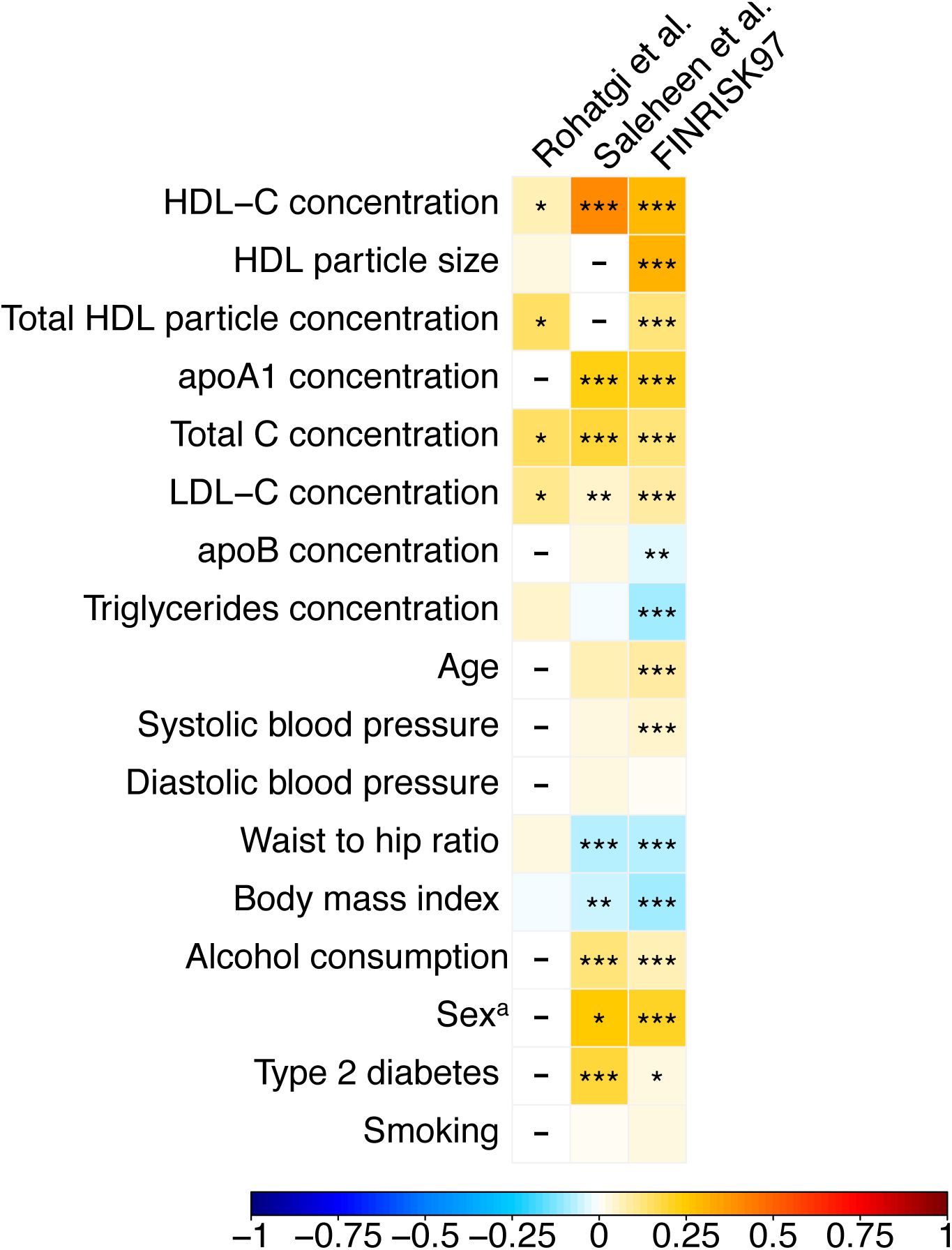
Associations of HDL-CEC with clinical and lipid parameters. In Rohatgi et al^11^ data are Spearman correlation coefficients and in Saleheen et al^10^ data were analysed by linear regression adjusted for age and sex. In the FINRISK97 data are Spearman correlation coefficients adjusted for age and sex; n = 7,370 (exact correlation coefficients and details in **Supplementary Table 2**). “-”in the figure denotes that a correlation coefficient was not available. ^a^association for female sex. ***P < 0.001, **P < 0.01, *P < 0.05.

Previously Rohatgi et al^11^ examined myocardial infarction and cardiovascular disease (CVD) in 2,416 individuals over a median follow-up of 9.4 years in a population-cohort with 30 and 172 endpoints, respectively, and Saleheen et al^10^ used a prospective nested case-control study with 1,745 patients with incident coronary heart disease (CHD) and 1,749 control participants. We chose these largest HDL-CEC endpoint studies as a reference. **Supplementary Table 2** presents comparisons for the associations of the *in vitro* measured HDL-CEC in cAMP-treated J774 cells with MI and CVD (Rohatgi et al^11^) and CHD (Saleheen et al^10^) outcomes with the current results for the NMR-based HDL-CEC estimates and CHD in the FINRISK97 cohort. All the main associations based on the *in vitro* estimates, including those with various adjustments, were replicated with the NMR-based estimates (see online Methods for Statistical analyses). Consistent with the findings by Rohatgi et al^11^ and Saleheen et al^10^ our independent large-scale study also supports the inverse association of HDL-CEC for cardiovascular outcomes. An additional consistent feature in the prior studies together with our data is that the associations of HDL-CEC with vascular outcomes are robust to multiple adjustments, including HDL-C, apoA1 and total HDL particle concentration (HDL-P).

Rohatgi et al^11^ also analysed the improvement in risk prediction after adding HDL-CEC with traditional risk factors in prediction models. This resulted in small improvements in all the risk-prediction indexes for the primary end point (consisting of atherosclerotic CVD), including changes in the C-statistic from 0.827 to 0.841 (P = 0.02), the integrated discrimination improvement index (IDI) of 0.02 (P < 0.001), and the net reclassification index (NRI) of 0.37 (95%CI, 0.18, 0.56). In our analyses with well-calibrated models (**Supplementary Fig. 4**), addition of HDL-CEC to traditional risk factors and HDL-C was also associated with small improvements in CHD prediction with improvements in the C-statistic (from 0.841 (95%CI, 0.829, 0.854) to 0.843 (95%CI, 0.830, 0.856); P = 0.02 by Student t-test for dependent samples: t.stat = 1.20, df = 7290, one-sided), IDI of 0.005 (95%CI, 0.001, 0.015), and NRI of 0.21 (95%CI, 0.05, 0.28). Similar improvements were observed for CVD prediction with C-statistic (from 0.837 (95%CI, 0.825, 0.848) to 0.838 (95%CI, 0.826, 0.849]; P = 0.01 by Student t-test for dependent samples: t.stat = 2.29, df = 7230, one-sided), IDI of 0.004 (95%CI, 0.001, 0.010), and NRI of 0.18 (95%CI, 0.04, 0.26). Hosmer-Lemeshow statistics for model calibration with CVD were X^2^ = 0.021, P = 1, df = 8 and X^2^ = 0.020, P = 1, df = 8 for models with and without HDL-CEC, respectively.

**Figure 4:**
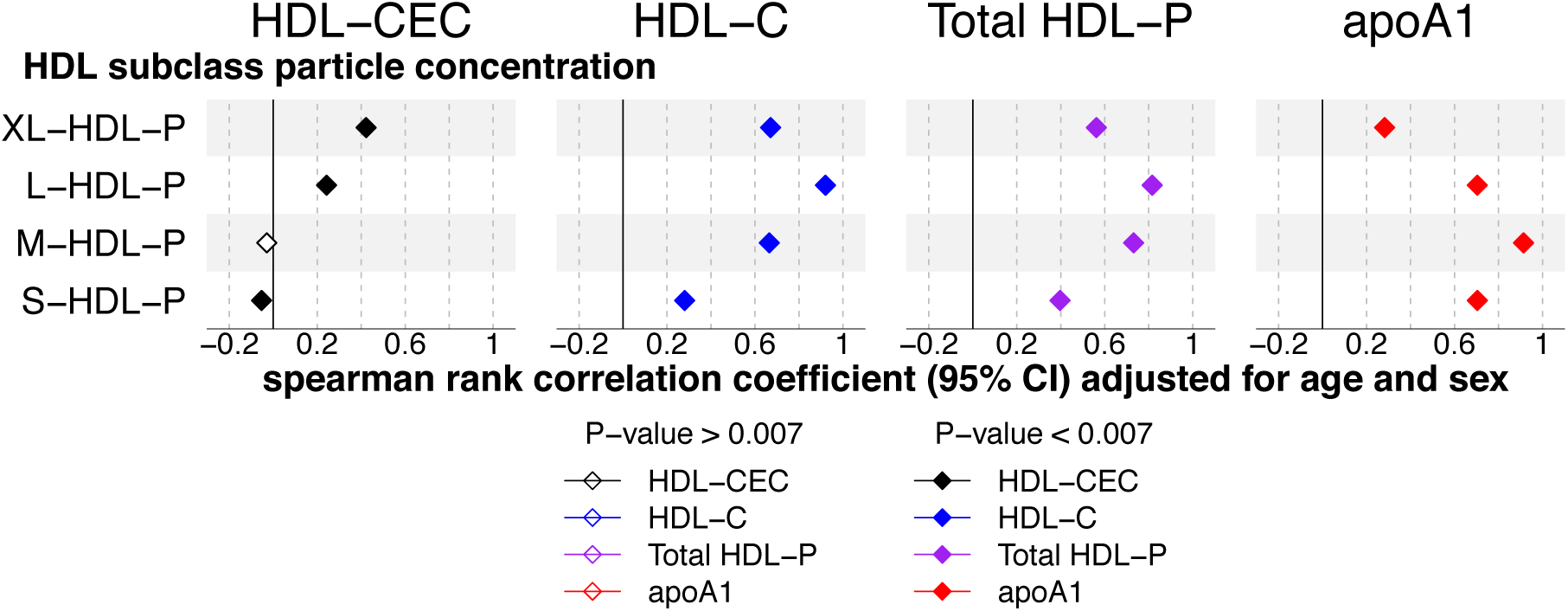
Associations of HDL-CEC and related measures with HDL subclass particle concentrations in the FINRISK97 cohort (n = 7,597). Data are Spearman correlation coefficients (95% CI) adjusted for age and sex (df = 7,595). HDL subclasses were measured by NMR spectroscopy and are defined by particle size as follows: XL-HDL; very large (average particle diameter 14.3 nm), L-HDL; large (12.1 nm), M-HDL; medium (10.9 nm) and S-HDL; small HDL (8.7 nm)^19^. P-value was adjusted for multiple testing by using principal component analysis (online Methods). HDL-CEC; NMR-based HDL-mediated cholesterol efflux estimate, HDL-C; high-density lipoprotein cholesterol concentration, apoAl; apolipoprotein Al concentration, HDL-P; total high-density lipoprotein particle concentration (a sum of the individual HDL subclass particle concentrations).

We also wanted to compare how the associations for our FINRISK97 cohort using the NMR-based HDL-CEC estimates compared with those from previous studies, as reported in a recent meta-analysis^14^. **Fig. 2** presents the association of HDL-CEC with cardiovascular risk in this study with comparable data from the meta-analysis^14^. The results in the FINRISK97 study were remarkably similar to the meta-analysis summary estimates. For example, the RR estimate for the highest vs lowest quartile of HDL-CEC was 0.69 (95%CI: 0.54, 0.87) **(Fig. 1)** in the FINRISK97 cohort, consistent with the meta-analysed RR estimate of 0.58 (0.39, 0.86)^14^. Similarly, the per 1-SD higher HDL-CEC RR estimate of 0.86 (0.79, 0.93) **(Supplementary Table 1**) in the FINRISK97 cohort was near-identical to the summary meta-analysis RR of 0.87 (0.73, 1.04)^14^. For both of these, there was no evidence of heterogeneity between the current HDL-CEC estimates and the pooled estimates from the meta-analysis (P value 0.46 and 0.91 for highest vs lowest and per-1-SD higher, respectively).

### Correlations of HDL-CEC with various anthropometric, lipoprotein and clinical variables

The cross-sectional correlations of NMR-based HDL-CEC estimates with various traits in the FINRISK97 cohort are presented in **Fig. 3** (exact correlation coefficients are given in **Supplementary Table 3)** together with the corresponding values from Rohatgi et al^11^ and Saleheen et al^10^. Analogous to the previous studies, we identified positive correlations of HDL-CEC with, for example, HDL-C, HDL particle size, age, blood pressure, alcohol consumption and female sex. Negative correlations of HDL-CEC were identified for apolipoprotein B, triglycerides, and measures of adiposity. The magnitudes of these correlations were weak, which corresponds with findings reported by Saleheen et al^10^. The correlations of HDL-CEC with HDL-C and HDL particle size were even weaker in Rohatgi et al^11^. While HDL-CEC was positively correlated with HDL-C and other HDL-related measures, the highest correlations of HDL-CEC with HDL-C across these 3 studies was 0.4 (**Supplementary Table 3**), supporting the view that HDL-CEC and HDL-C contain largely independent information on HDL metabolism.

### Association of HDL-CEC with HDL subclasses

Serum NMR spectroscopy enables the extensive quantification of lipoprotein subclasses, including particle concentrations for four HDL subclasses^18,19^. These data were not available in prior studies including Rohatgi et al^11^ or Saleheen et al^10^. **Figure 4** shows correlations between the particle concentrations of four HDL subclasses (extra large, large, medium and small) and the NMR-based HDL-CEC estimates, HDL-C, apoA1 and total HDL-P in FINRISK97. In contrast to HDL-CEC, the other HDL-related measures were positively correlated with all the HDL subclass particle concentrations. However, HDL-CEC correlated only with the two largest HDL particle categories. More detailed association analysis between the NMR-based HDL-CEC estimates and HDL subclass lipids are shown in **Supplementary Fig. 5**.

## DISCUSSION

We developed a high-throughput method to estimate HDL-CEC directly from serum samples using NMR spectroscopy. The method is based on HDL-CEC estimates from *in vitro* measurements in cAMP-treated J774 cells, the most common technique to analyse HDL-CEC in cardiometabolic research. The NMR-based HDL-CEC estimates appear to capture the same key aspects as the *in vitro* HDL-CEC with respect to associations with various anthropometric, lipoprotein and clinical variables as well as, importantly, with cardiovascular outcomes. We applied the new NMR-based method to estimate HDL-CEC in a large-scale epidemiological study, the FINRISK97 cohort of ∼7,200 individuals, among whom 574 CHD and 789 CVD incident events occurred during 15 years of follow-up. This is currently the largest epidemiological study of HDL-CEC, the results of which also show independent inverse associations of HDL-CEC with risk of CHD and CVD lending support to a potential atheroprotective role of HDL function.

The associations of NMR-based HDL-CEC with risk of cardiovascular outcomes were in keeping with summary estimates of a recent meta-analysis investigating the associations of cellular *in vitro* HDL-CEC estimates with cardiovascular outcomes^14^. A key finding with the NMR-based HDL-CEC, corroborating the earlier findings with the cellular *in vitro* HDL-CEC in the existing largest outcome studies^10,11^ was its association with CHD being independent of other HDL-related measures, including HDL-C, apoA1 and HDL-P. Therefore, the NMR-based HDL-CEC estimates recapitulate the characteristics of cellular *in vitro* HDL-CEC estimates and account for independent information on CHD risk, not captured by other HDL-related measures.

Cardiovascular risk prediction is rarely improved by new biomarkers^21^. Currently, the ability of HDL-CEC to improve CVD risk prediction beyond traditional risk factors has been investigated in two studies, both describing moderate increases in NRI: 38%^16^ and 22%^11^, but relatively small increases in C-statistics. Accordingly, we found an increase in NRI of 21% and small increases in the C-statistics. A small increase in C-statistic is expected, since this metric is insensitive in model comparisons, when good predictors are already present in the reference model^22^.

HDL-CEC is a functional measure related to multiple characteristics of HDL particles, and therefore some level of correlation with other HDL-related measures would be expected. In the FINRISK97 cohort, relatively weak correlations were observed, with the largest being between the NMR-based HDL-CEC estimate and mean HDL-particle size (correlation coefficient 0.31). In Saleheen et al^10^ the highest correlation was 0.4 between the cellular *in vitro* HDL-CEC estimate and HDL-C. In the GRAPHIC cohort with HDL-CEC data in 1,988 individuals, a correlation of 0.62 between HDL-C and HDL-CEC was found^17^ and in the JUPITER trial data the highest correlation was 0.48 between HDL-CEC and apoA1^13^. Together with the independent associations of HDL-CEC with risk of vascular disease, this points towards HDL-CEC containing independent information on HDL metabolism and reverse cholesterol transport.

The present study is the first large-scale study investigating the association of HDL-CEC with HDL subclass measures. The HDL-CEC estimates associated with very large and large, but not with medium and small, HDL subclass particle concentrations. These findings match previous small-scale studies^23,24^ and the predominance of the associations with larger HDL particles is most likely due to these particles having a larger receiving area for the diffusing cholesterol molecules and thereby more effective mediation of diffusion than smaller particles^9^. This is particularly relevant here since diffusion is thought to be the dominating mechanism for the cholesterol efflux in radioactive cholesterol labelled cAMP-treated J774 cells (see online Methods).

The new cost-effective NMR-based method presented here to estimate HDL-CEC directly from serum samples is designed to correspond to the most common assay to analyse HDL-CEC, i.e., using cAMP-treated J774 murine macrophages with radiolabelled cholesterol^10,12–14^. This latter methodology was used by Saleheen et al^14^, a large-scale outcome study we compared our current results with. The excellent correspondence of our findings (using the NMR-based HDL-CEC estimates) to those by Saleheen et al^10^ (using cAMP-treated J774 murine macrophages), serves to corroborate our approach and is in part expected, given that our NMR assay was developed from the same in vitro assay. In contrast, the other large-scale outcome study by Rohatgi et al^11^ applied a less common, fluorescence-labelled cholesterol method in a similar cell model^11,15^. However, the correlation between HDL-CEC estimates from these two cellular *in vitro* assays is quite high^11,15^ and it is therefore expected that the NMR-based HDL-CEC results in the FINRISK97 cohort would also match results by Rohatgi *et al*^15^. Our data are also consistent with the pooled estimate of a recent meta-analysis that summarises the associations of *in vitro* HDL-CEC estimates with cardiovascular outcomes^14^, which serves to further corroborate our NMR-quantified HDL-CEC.

Multiple recent drug trials have indicated that increasing circulating HDL-C concentrations does not lead to a reduction in cardiovascular disease^6^ Mendelian randomization studies also fail to support HDL-C having a causal role in cardiovascular diseases^4^. We should therefore remain sceptical about the potential causality of HDL-CEC. Nevertheless, the new cost-effective NMR-based method to estimate HDL-CEC could be advantageous in widening the research of cholesterol efflux to large population-based cohorts and drug trials and to expedite appropriately powered studies in relation to multiple cardiovascular and metabolic outcomes. Large-scale cohorts with HDL-CEC estimates are needed to investigate and replicate the associations of HDL-CEC with clinical outcomes, and more importantly, to study the genetic determinants of cholesterol efflux in order to perform Mendelian randomization analyses for potential causality. We propose the new NMR-based method as a pragmatic alternative for HDL-CEC estimates from *in vitro* measurements in cAMP-treated J774 cells, particularly in large-scale epidemiology and genetics. This method appears to have great potential to lower the experimental costs related to HDL-CEC measurements and concomitantly speed-up collection of the extensive epidemiological evidence-base necessary to ascertain whether this functional HDL-phenotype is causal for vascular disease and thus, whether it provides an opportunity for translational applications.

## ONLINE METHODS

### Training data

Random blood samples were collected during 2016 from Finnish Red Cross blood service in accordance with the ethical guidelines required by the Helsinki Declaration. Serum was obtained by centrifugation at 1500 x g for 15min at ambient temperature and stored at −80°C. HDL-CEC was measured using cAMP-treated J774 cells within a year of sample collection. The same serum samples were also analysed by proton NMR spectroscopy within the same time frame. The complete training data set obtained comprised 199 individuals with the serum NMR spectra and the corresponding cellular *in vitro* HDL-CEC estimates. Bayesian regression modelling was applied to link the NMR spectra to the HDL-CEC estimates^25^; the correspondence between the NMR-based and the cellular *in vitro* HDL-CEC estimates are shown in **Supplementary Fig. 2.** Characteristics of the individuals in the training data set are given in **Supplementary Table 4**.

### Cellular *in vitro* HDL-CEC measurements

Efflux experiments were performed for the apoB-depleted serum samples in the training data set. ApoB-containing lipoprotein precipitation was performed by polyethylene glycol (PEG) to obtain an HDL fraction for cholesterol efflux studies^10,12,13^. Briefly, 40 parts of precipitation-solution (20% PEG in 200 mM glycine buffer, pH 7.4) were added to 100 parts of serum and mixed. PEG and glycine were purchased from Fisher Scientific. After 20 min of incubation, the samples were centrifuged (10 000 x g, 40 min, 4°C) and the supernatant containing the HDL fraction was recovered. A day before the efflux experiments, apoB-depleted serum samples were diluted to 5.6% in MEM-HEPES (containing 10 mM HEPES in MEM; both purchased from Sigma).

The cholesterol efflux capacity of apoB-depleted serum (HDL-CEC) was measured by commonly used radiolabelled cholesterol assay using cAMP-treated J774 murine macrophages^10,12–14^. The efflux experiments were performed with J774 cells at 37°C in humidified atmosphere of 5% CO_2_ and 95% air. Briefly, cells were plated (7 x 10^5^ cells/well) in 96-well plates, treated with cAMP and labelled with radioactive cholesterol in serum free medium for overnight. Labelling medium contained 0.3 mM cAMP (8-(4-Clorophenylthio)-cyclic AMP, Sigma), 4 μCi/mL [1,2-^3^H(N)]- cholesterol (Perkin Elmer), 2 mM L-Glutamine (Sigma) and 1% (v/v) penicillin–streptomycin (Sigma) in DMEM (Sigma). Next day, cells were washed with MEM-HEPES and efflux assays were performed. To start the efflux period, 0.3 mM cAMP in MEM-HEPES was added first on the cells, following addition of apoB-depleted serum samples. During the efflux period, 2.8% apoB-depleted serum samples were incubated with the labelled cells for 6 hour in MEM-HEPES containing 0.15 mM cAMP. Thereafter medium was collected and cell lipids extracted by 2-propanol. Aliquots of the medium and cell extracts were counted for radioactivity by β-counter and cholesterol efflux was expressed as percentage of radioactive cholesterol released in the medium from total radioactive cholesterol present in the well. For each serum sample, assays were done as parallel measurements and control serum was included for all plates. Intra-assay and inter-assay CV% were 3.6% and 6.1%, respectively.

The efflux pathways in radioactive cholesterol labelled cAMP-treated J774 cells are ABCA1, ABCG1, SR-B1, passive diffusion and possible undiscovered pathways^12^. In this cell model, the majority of cholesterol efflux seems to be mediated through diffusion, either passively or by ABCG1 and SR-B1, and possible unknown pathways^12^. However, the role of ABCA1 pathway varies depending on cAMP-treatment conditions, since this efflux receptor is inducible by cAMP^12,26^. Therefore, we determined the percentage of efflux mediated trough ABCA1 pathway by commonly used robust assay: ABCA1-efflux = efflux % with cAMP-treatment minus efflux % without cAMP-treatment^27^. For this measurement, we performed parallel efflux-analysis without cAMP-treatment simultaneously in same efflux-assay with *in vitro* HDL-CEC measurement. A median of 7% (interquartile range: 5%-10%, n = 199) of the total cholesterol efflux was mediated by robust ABCA1 pathway, indicating that majority of the efflux was mediated through diffusion (either passively or facilitated by ABCG1^9^ and SR-B1^9^) or undiscovered pathways.

### NMR spectroscopy and sample preparation

A high-throughput NMR spectroscopy platform with an optimised measurement protocol was used to provide quantitative information on the multiple molecular constituents of serum. The experimental details have been previously published^28^.

### Lipoprotein quantification and HDL-CEC modelling from the NMR spectra

Lipoprotein profiling by proton NMR spectroscopy is currently well established and was done as previously described^18,19,28^. A computationally more efficient modification of the Bayesian approach than we presented previously^25^ was applied. The lipoprotein subclasses in NMR spectroscopy are defined by particle size: potential chylomicrons and the largest very-low-density lipoprotein particles (XXL-VLDL; average particle diameter *≥*75 nm); five different VLDL subclasses, i.e. very large (average particle diameter 64.0 nm), large (53.6 nm), medium (44.5 nm), small (36.8 nm) and very small VLDL (31.3 nm); intermediate-density lipoprotein (IDL; 28.6 nm); and three LDL subclasses, i.e. large (25.5 nm), medium (23.0 nm) and small LDL (18.7 nm). The four size-specific HDL subclasses are very large (14.3 nm), large (12.1 nm), medium (10.9 nm) and small HDL (8.7 nm)^18,19^.

Similarly, selected regions of the NMR spectra (containing the main resonances of the lipoprotein lipid constituents) were linked, via a linear Bayesian model^25,28^, to the independently assayed HDL-CEC estimates from the cellular *in vitro* measurements in the training data set. A good correspondence between the NMR-based and the *in vitro* determined HDL-CEC estimates was established with a cross-validation R^2^ = 0.83; **Supplementary Fig. 2**. The analytical correspondence is similar to those between NMR-based lipoprotein lipid measures and the corresponding measures from traditional clinical chemistry assays^19,25^. In addition, similar correspondence has also been indicated between NMR-based and clinical biochemistry for eight routinely measured traits^29^. This established linear Bayesian model (together with the lipoprotein profiling) was applied in the FINRISK97 cohort for all the 7,603 serum NMR spectra to estimate corresponding HDL-CEC values; the distribution of these HDL-CEC estimates is illustrated in **Supplementary Fig. 6**.

### Epidemiological study population and statistical analyses

The FINRISK97 survey was carried out to monitor the health of the Finnish population among persons aged 25-74 at recruitment^30^. The study was conducted in 5 study areas across Finland, recruiting a total of 8444 persons. The Ethics Committee of the National Public Health Institute, Helsinki, Finland has approved the study in accordance with Declaration of Helsinki and written informed consent has been obtained from all participants. Serum samples were collected at semi-fasting state (median fasting time 5h; interquartile range: 4-6 hour).

For parametric statistics, outliers for each quantitative measure were defined as values that lie over 4 times higher or lower than the interquartile range. In the case of HDL-CEC, 74 outliers were removed from the statistical analyses. In addition, for the analysis of the association of NMR-based HDL-CEC estimates with incident CHD events, individuals with prevalent CHD events (n=203) and missing data and outliers in study variables (n = 110) were removed leaving complete data for 7,290 individuals and 574 incident CHD events (defined as fatal or nonfatal myocardial infraction, cardiac revascularization, or unstable angina) for the analysis. For the analysis of the association of NMR-based HDL-CEC estimates with incident CVD events (defined as fatal or nonfatal myocardial infraction, ischemic stroke, cardiac revascularization, or unstable angina), individuals with prevalent CVD events (n = 263) and missing data and outliers in study variables (n = 109) were removed leaving complete data of 7,231 individuals and 789 incident CVD events for the analysis. The median follow up time was 14.8 years. Baseline characteristics are presented in **Supplementary Table 4**. For non-parametric statistics (Spearman’s rank correlation), all participants with complete data were used.

Statistical analyses were performed by R statistical software version 3.3.3. The survival analysis was performed by R packages survival, survminer and SurvMisc. The associations of HDL-CEC estimates with incident CHD events were analysed by Cox proportional hazard regression models. For these analyses skewed variables were transformed for normality: triglyceride concentrations and BMI were log-transformed and alcohol intake log1p-transformed (log_e_ (1+x)). The fully adjusted model included age, sex, geographical region, diabetes, mean arterial blood pressure, blood pressure treatment, smoking, log1p alcohol intake, log BMI, waist to hip ratio, LDL cholesterol, log triglycerides, lipid lowering treatment and concentrations of HDL cholesterol, HDL particle and apolipoprotein A1. A key aim was to see if the NMR-based HDL-CEC estimates would show similar associations to previous large-scale epidemiological studies. Therefore, we performed analyses as similar to those used by Rohatgi et al^11^ and Saleheen et al^10^, including using the same statistical adjustments, and calculated the hazard ratios for incident CHD events for tertiles and quartiles using the bottom one as the reference. In addition, analyses were performed per 1-SD higher HDL-CEC (see **Supplementary Table 2**).

The ability of NMR-based HDL-CEC estimates to improve CHD risk prediction beyond traditional risk factors and HDL-C was assed in well-calibrated models (**Supplementary Fig. 4**) by using C-statistics, net reclassification index (NRI) and integrated-discrimination-improvement index (IDI). To evaluate model calibration, predicted and observed risk was assessed by R package rms. Hosmer-Lemeshow test was calculated by R package MKmisc. C-statistics was calculated by R package survcomp and NRI & IDI by the R package survIDINRI.

Associations between variables were assessed by partial correlation (Spearman) adjusted for age and sex. Partial correlation coefficients (95% CI and p-value) were calculated by R package RVAideMemoire using 500 replications to compute 95% confidence intervals by bootstrapping. Two-sided P-value was computed through the asymptotic t approximation. Due to overall cross-correlation nature of HDL subclass data, we made corrections for multiple tests when analysing associations of HDL-related measured with HDL subclass data by evaluating the number of independent tests using principal component analysis. Seven principal components were able to explain 99% of variation in the FINRISK97 HDL subclass data. Due to this, P < 0.007 (0.05/7) was used to denote statistical significance in these analyses.

## Availability of data

The FINRISK97 cohort is deposited in the biobank of the Institute of Health and Welfare, Finland and can be applied through the biobank.

## Code availability

Code used to estimate NMR-based HDL-CEC from NMR spectra is owned by Nightingale Health Ltd. and therefore is not available for public use.

## Author contributions

S.K. and M.A.-K. conceived and designed the study, interpreted the results and wrote the manuscript. S.K. performed the cellular experiments and the statistical analyses and A.J.K. the spectral modelling and bioinformatics. M.K. and M.T. prepared the samples and performed the NMR experiments. M.V.H., P.O. and J.K. interpreted the results and edited the manuscript. M.P. and V.S. provided samples and the phenotype data of FINRISK97. All authors discussed the results and approved the final version of the manuscript. M.A.-K. supervised the study.

